# The dynamic core microbiome: Structure, dynamics and stability

**DOI:** 10.1101/137885

**Authors:** Johannes R. Björk, Robert B. O’Hara, Marta Ribes, Rafel Coma, José M. Montoya

## Abstract

The long-term stability of microbiomes is crucial as the persistent occurrence of beneficial microbes and their associated functions ensure host health and well-being. Microbiomes are highly diverse and dynamic, making them challenging to understand. Because many natural systems work as temporal networks, we present an approach that allows identifying meaningful ecological patterns within complex microbiomes: the dynamic core microbiome. On the basis of six marine sponge species sampled monthly over three years, we study the structure, dynamics and stability of their microbiomes. What emerge for each microbiome is a negative relationship between temporal variability and mean abundance. The notion of the dynamic core microbiome allowed us to determine a relevant functional attribute of the microbiome: temporal stability is not determined by the diversity of a host’s microbial assemblages, but rather by the density of those microbes that conform its core microbiome. The core microbial interaction network consisted of complementary members interacting weakly with dominance of comensal and amensal interactions that suggests self-regulation as a key determinant of the temporal stability of the microbiome. These interactions have likely coevolved to maintain host functionality and fitness over ecological, and even evolutionary time scales.

## Introduction

Microbes form intricate relationships with most animals and plants, with symbiosis postulated as one of the driving forces behind diversifications across the tree of life ([1]). Research on host-microbe symbioses are typically restricted to highly specialized reciprocal interactions with one or a few microbes interacting with a single host, resulting in mutual benefits for both parties ([2, 3]). However, more diverse and complex host-associated microbial communities (hereafter *microbiomes*) are increasingly found in different plant and animal species ([1]). This poses a challenge because the pairwise specificity, coevolution and reciprocity of host-microbe interactions might not explain the structure, dynamics and functioning of microbiomes. The mere existence of multiple microbes interacting with a host suggests that interactions among microbes might also be an important driver regulating the overall abundance and composition of microbiomes and their associated ecosystem stability and functions.

The diversity, complexity and highly dynamic nature of microbiomes makes them challenging to understand. We thus require approaches that embrace the complexity and dynamics but still allow identifying meaningful ecological patterns within and across microbiomes. The quest for core microbiomes is a promising avenue. A core microbiome is typically defined cross-sectionally, rather than longitudinally ([4, 5, 6, 7, 8, 9, 10]), thus failing to capture temporal dynamics. However, a core microbiome characterized as a set of microbes consistently present over long periods of time ([11, 12, 13]), is more likely to be important for the development, health and functioning of its host; for example, several aspects of human health, including autoimmune disorders ([14, 15]), diabetes ([16]) and obesity ([17, 5]) can be linked to severe shifts in the gut microbiome. Whether these disorders emerge as a consequence of perturbed core microbiomes (defined longitudinally) still remains to be seen. However, arguably, the long-term stability of the core microbiome is likely critical as the persistent occurrence of beneficial microbes and their associated ecosystem functions ensure host health and well-being ([18, 19, 20, 21, 22]).

Despite the recent realizations that complex microbiomes pervade the tree of life, little is known about microbiome dynamics beyond humans. Here we study the structure, dynamics and stability of microbiomes from six coexisting marine sponges (*Porifera*) belonging to different orders that were sampled over 36 consecutive months.

Sponges are keystone species in marine coastal areas due to their filter-feeding activities: they regulate primary and secondary production by transferring energy between the pelagic and benthic zones ([23, 24]). Despite their constant influx of water, they maintain highly diverse, yet specific microbiomes with very little intraspecific variation ([25]). As *Porifera* is a sister-group to all other multicellular animals ([26]), their association with microbes is likely the oldest extant form of an host animal-microbe symbiosis ([27, 28, 29]).

We analyzed sponge species corresponding to two different groups that differ markedly in numerous traits illustrating their dependence upon their association with microbes. This classification is based on the diversity and abundance of microbes they harbor–High and Low Microbial Abundance (HMA and LMA) sponges. The classification pervades sponge host morphology and physiology; LMA hosts have an interior architecture fitted for pumping large volumes of water, whereas HMA hosts are morphologically adapted to harbor denser microbial assemblages within their tissue ([30, 31]). As a result, LMA hosts are more dependent on nutrient uptake from the water column ([32, 30, 31, 33, 34]) compared to HMA hosts that rely more heavily on nutrients produced by their microbial symbionts ([33, 34, 35, 36, 37, 38]). These two sets of hosts provide an ideal system from which we can generalize whether the structure and temporal dynamics of complex microbiomes differ across hosts with different eco-evolutionary characteristics and lifestyles.

Our general aim is to understand the temporal dynamics of complex microbiomes. More specifically, we try to answer: (i) What is the diversity and community structure of each core microbiome and how does it differ from that of the many transient taxa passing through the host? (ii) What are the temporal dynamics and stability of each microbial taxon and their aggregated effect on the core microbiome? And (iii) what are the likely ecological processes that underpin the observed temporal dynamics and stability? We expect the answer to each of these questions to differ across hosts with different lifestyles (i.e., HMA vs. LMA), reflecting their different dependency on their respective microbiomes. In particular, we expect the core microbiomes harbored by HMA hosts to be more diverse, compositionally similar and less variable over time.

## Results

We analyzed six microbiomes–Three belonging to host species classified as HMA (*Agelas oroides; Chondrosia reniformis; and Petrosia ficiformis*), and three from hosts classified as LMA (*Axinella damicornis; Dysidea avara; and Crambe crambe*) ([39, 40])

### Opportunistic taxa dominate the sponge microbiome

To better understand the commonness and rarity of the taxa occurring throughout the time series, we divided each microbiome into three different temporal assemblages based on the persistence of individual taxa over the 36 consecutive months. Core microbiomes were defined as those taxa that were present in more than (or equal to) 70% of each time series (i.e., persisting ≥ 26 months), whereas opportunistic assemblages comprised taxa present in less than (or equal to) 30% of each time series (i.e., persisting ≤ 11 months). Intermediately persistent taxa (i.e., those persisting between 12 and 25 months) formed transient assemblages. This resulted in three temporal assemblages each spanning 36 months, with individual taxa not restricted to consecutively occur over the course of the time series. While this categorization is rather arbitrary, it agrees with previous classifications based on temporal occurrences ([41, 42]). Grime (*1998*) for example, suggested a similar approach (the mass ratio hypothesis) to better understand the relationship between plant diversity and ecosystem properties, dividing species into three categories–dominants, subordinates and transients–reflecting their contribution to ecosystem biomass, stability and functioning.

As typically observed for macroecological systems (see e.g., [43]), we found that opportunistic assemblages made up the bulk of the total diversity (i.e., species richness) that resulted from the aggregation of all taxa observed throughout the time series with the core microbiomes only representing a small fraction (1.24%) of this diversity (Table 1). Core taxa still represented an important fraction of the monthly diversity in some of the hosts; Compared to LMA hosts that on a monthly basis harbored 19 to 25 times more opportunistic taxa, core taxa were only 4 to 5 times less numerous in HMA hosts (Table 1). As common species are increasingly recognized as key contributors to ecosystem functions ([44, 45, 46]), the observed difference in the number of common taxa may mirror variation in the aforementioned symbiont dependency among HMA and LMA host species, respectively.

**Table 1:**
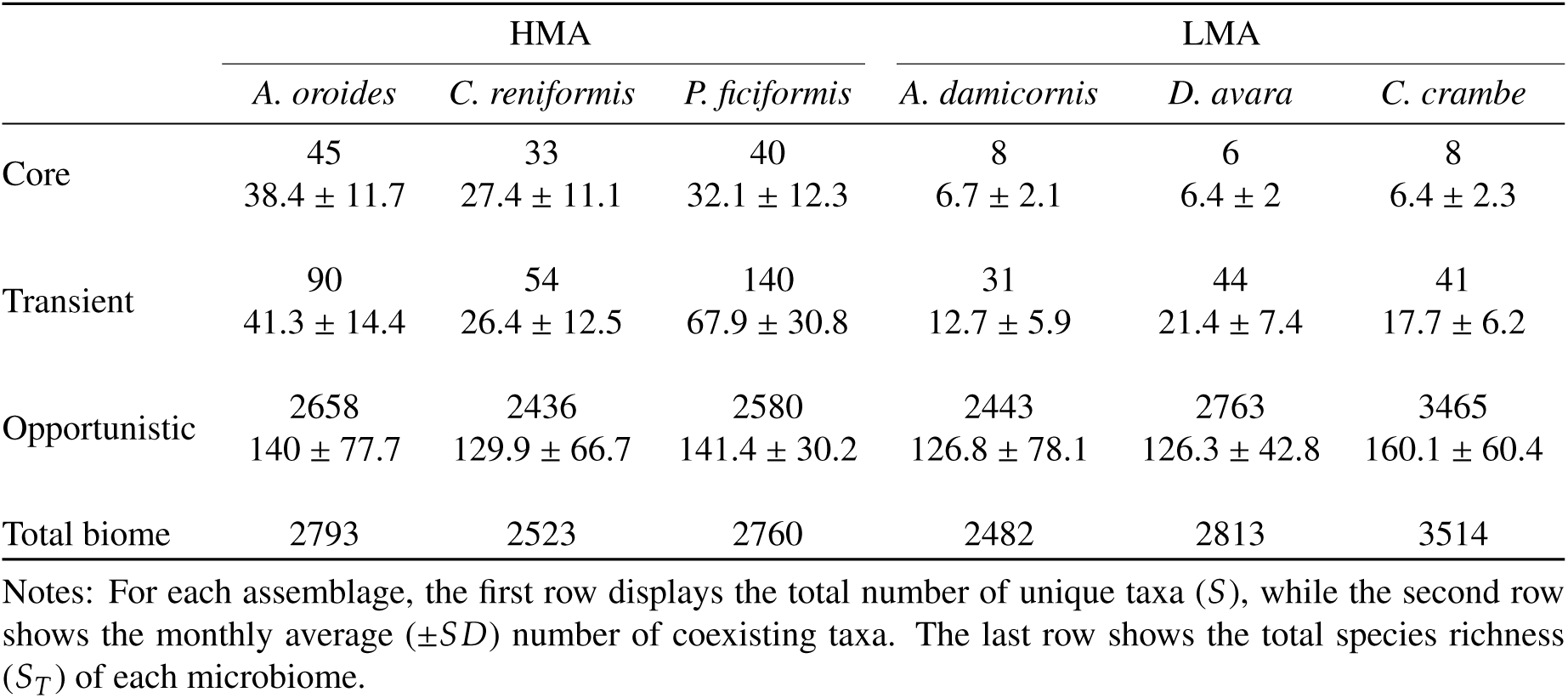
Microbiome diversity by temporal assemblage

|  | HMA |  |  | LMA |  |  |
| --- | --- | --- | --- | --- | --- | --- |
|  | *A. oroides* | *C. reniformis* | *P. ficiformis* | *A. damicornis* | *D. avara* | *C. crambe* |
| Core | 45<br>38.4 ± 11.7 | 33<br>27.4 ± 11.1 | 40<br>32.1 ± 12.3 | 8<br>6.7 ± 2.1 | 6<br>6.4 ± 2 | 8<br>6.4 ± 2.3 |
| Transient | 90<br>41.3 ± 14.4 | 54<br>26.4 ± 12.5 | 140<br>67.9 ± 30.8 | 31<br>12.7 ± 5.9 | 44<br>21.4 ± 7.4 | 41<br>17.7 ± 6.2 |
| Opportunistic | 2658<br>140 ± 77.7 | 2436<br>129.9 ± 66.7 | 2580<br>141.4 ± 30.2 | 2443<br>126.8 ± 78.1 | 2763<br>126.3 ± 42.8 | 3465<br>160.1 ± 60.4 |
| Total biome | 2793 | 2523 | 2760 | 2482 | 2813 | 3514 |

Notes: For each assemblage, the first row displays the total number of unique taxa ( $S$ ), while the second row shows the monthly average ( $\pm SD$ ) number of coexisting taxa. The last row shows the total species richness ( $S_T$ ) of each microbiome.

### Core microbiomes are mainly driven by changes in abundance

Temporal turnover is an intrinsic property of our definition of core microbiomes, transient and opportunistic assemblages; for example, as opportunistic taxa persist less than 30% of the total time series, these assemblages are bound to heavily fluctuate in microbial composition. Nevertheless, in order to quantify the temporal turnover of each assemblage, we applied a measure that disentangles the two additive determinants of temporal turnover: change in species composition and change in total abundance ([47]). We found that core microbiomes were mainly driven by changes in abundance, whereas transient and opportunistic assemblages were governed by changes in species composition (Figures S1). Due to this large turnover, i.e., new taxa rapidly replace extinct taxa, the opportunistic assemblages displayed an overall higher average evenness (0.83±0.17), measured as Pielou’s J over time, compared to the core microbiomes (0.64±0.25).

### Microbiome composition and specificity differ across host species and lifestyles

Microbiome composition varied markedly across host species (Figure 1; Phylogenetic composition, Figure S2), with nearly no overlap amongst core microbiomes (Figure 2). The observed community structure was different from the expectation generated by a null-model (Figure S3; See *Supplementary material* for more information). In addition, HMA and LMA host species displayed very different taxonomic profiles (Figure 3). HMA hosts harbored three dominant phyla that accounted for roughly half of their diversity: *Chloroflexi; Actinobacteria; and Acidobacteria*. Despite the dominance of these three phyla, each HMA host still kept a unique taxonomic fingerprint by harboring other phyla, such as *Gemmatimonadetes, Nitrospira*, and *Bacteroidetes*. The core microbiomes of LMA hosts, on the other hand, were largely dominated by taxa from a single phylum–*Proteobacteria*.

**Figure 1:**
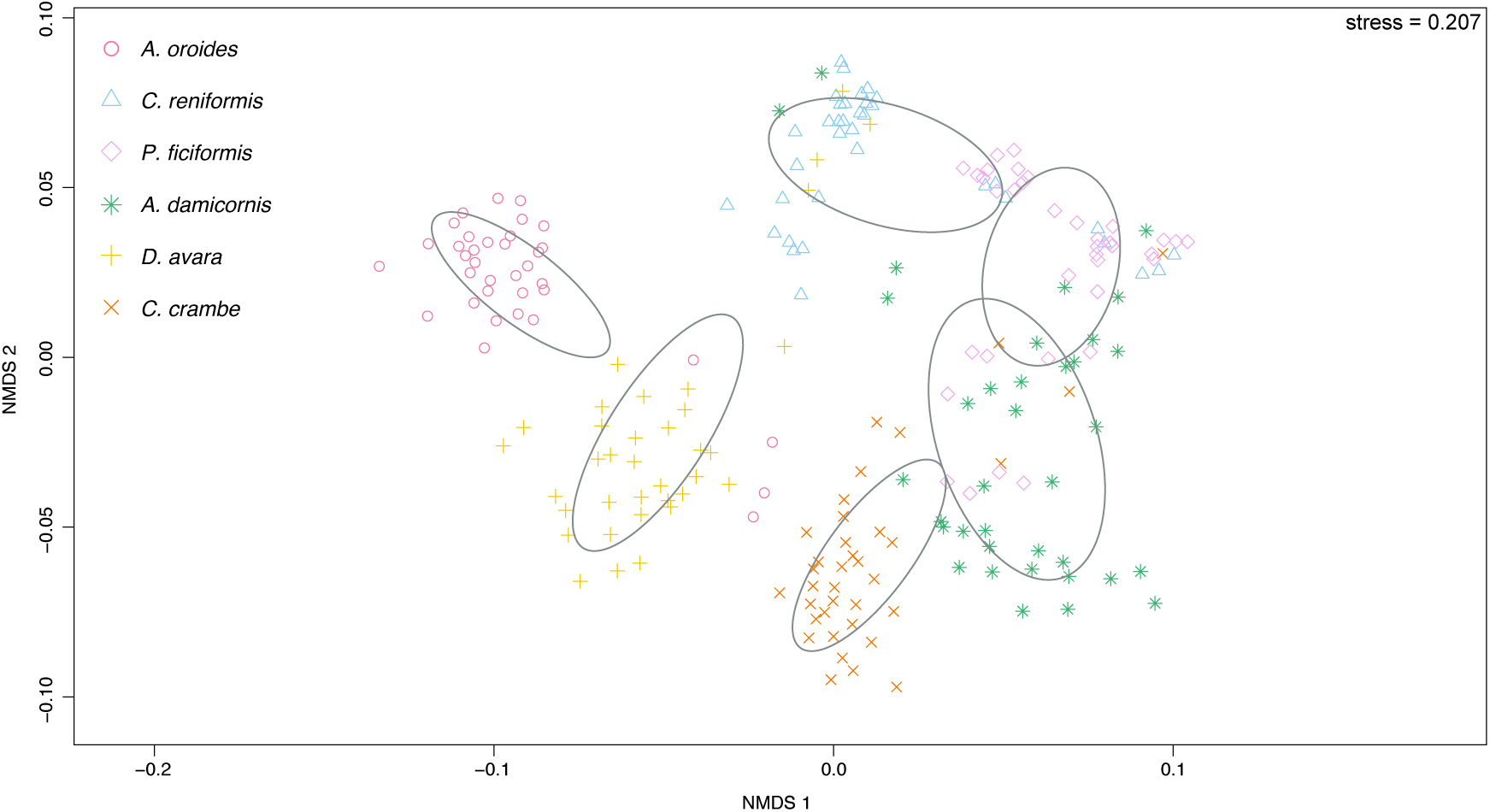
Microbiome compositional similarity across host species. Non-metric multidimensional scaling (NMDS) calculated on Jaccard distances among all 36 monthly samples for each host species (n=36x6=216). Colors and shapes denote all monthly samples from a given host species surrounded by an ellipse showing the 95% confidence interval. Red circles *A. oroides*; blue triangles *C. reniformis*; pink diamonds *P. ficiformis*; green stars *A. damicornis*; yellow crosses (+) *D. avara*; and orange crosses (⇥) *C. crambe*. ANOSIM: R=0.767, P<0.001

**Figure 2:**
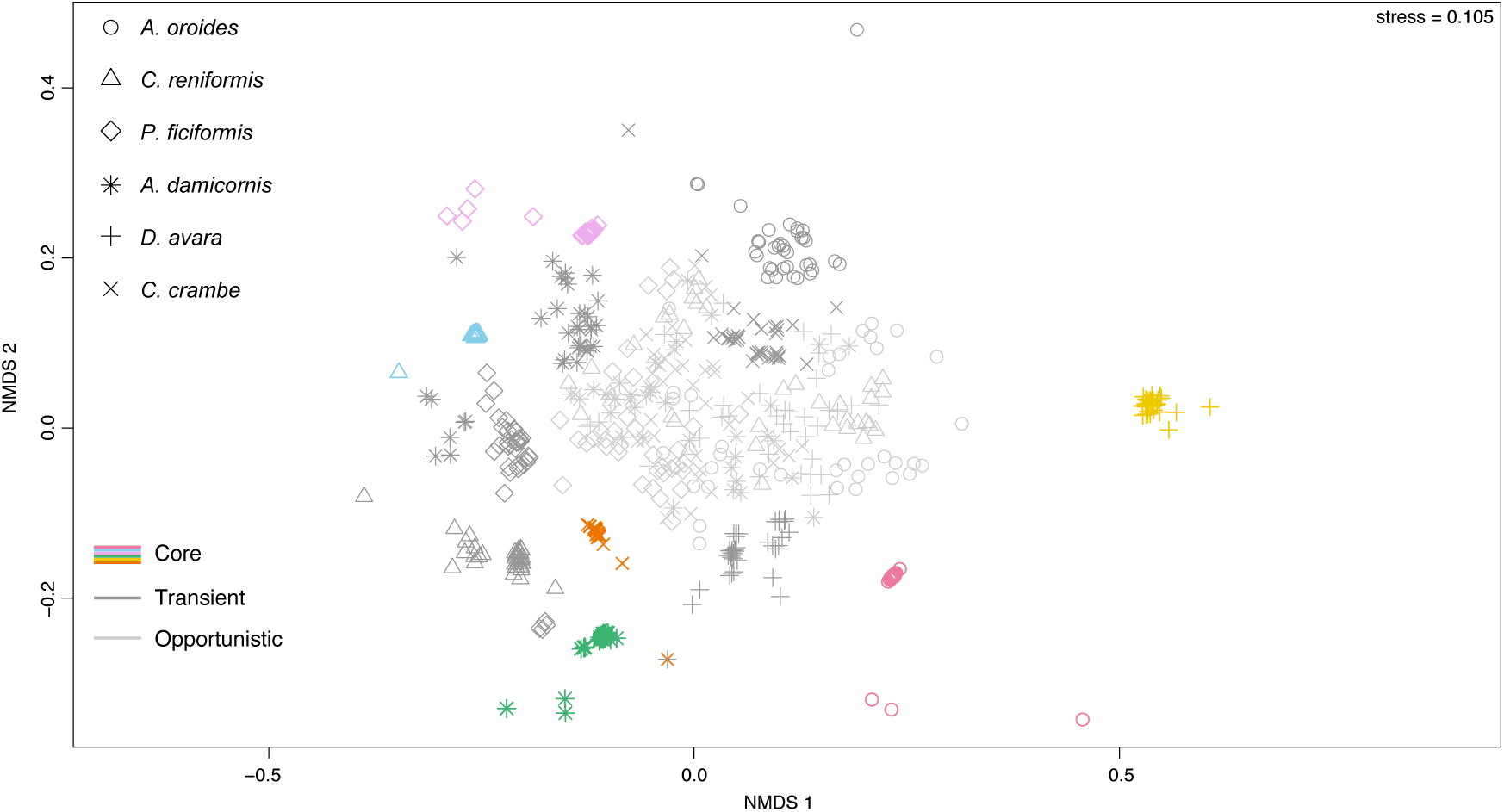
Microbiome compositional similarity across assemblages and host species. Non-metric multidimensional scaling (NMDS) calculated on Jaccard distances among all 36 monthly samples for each host and assemblage (n=36x6x3=648). Colors and shapes denote all monthly samples from a given host and assemblage, respectively. Different shapes for each host species: circles *A. oroides*; triangles *C. reniformis*; diamonds *P. ficiformis*; stars *A. damicornis*; crosses (+) *D. avara*; crosses (⇥) *C. crambe*. Different colors for each core microbiome: red *A. oroides*; blue *C. reniformis*; pink *P. ficiformis*; green *A. damicornis*; yellow *D. avara*; and orange: *C. crambe*. Dark gray denotes transient and light gray opportunistic assemblages, respectively.

**Figure 3:**
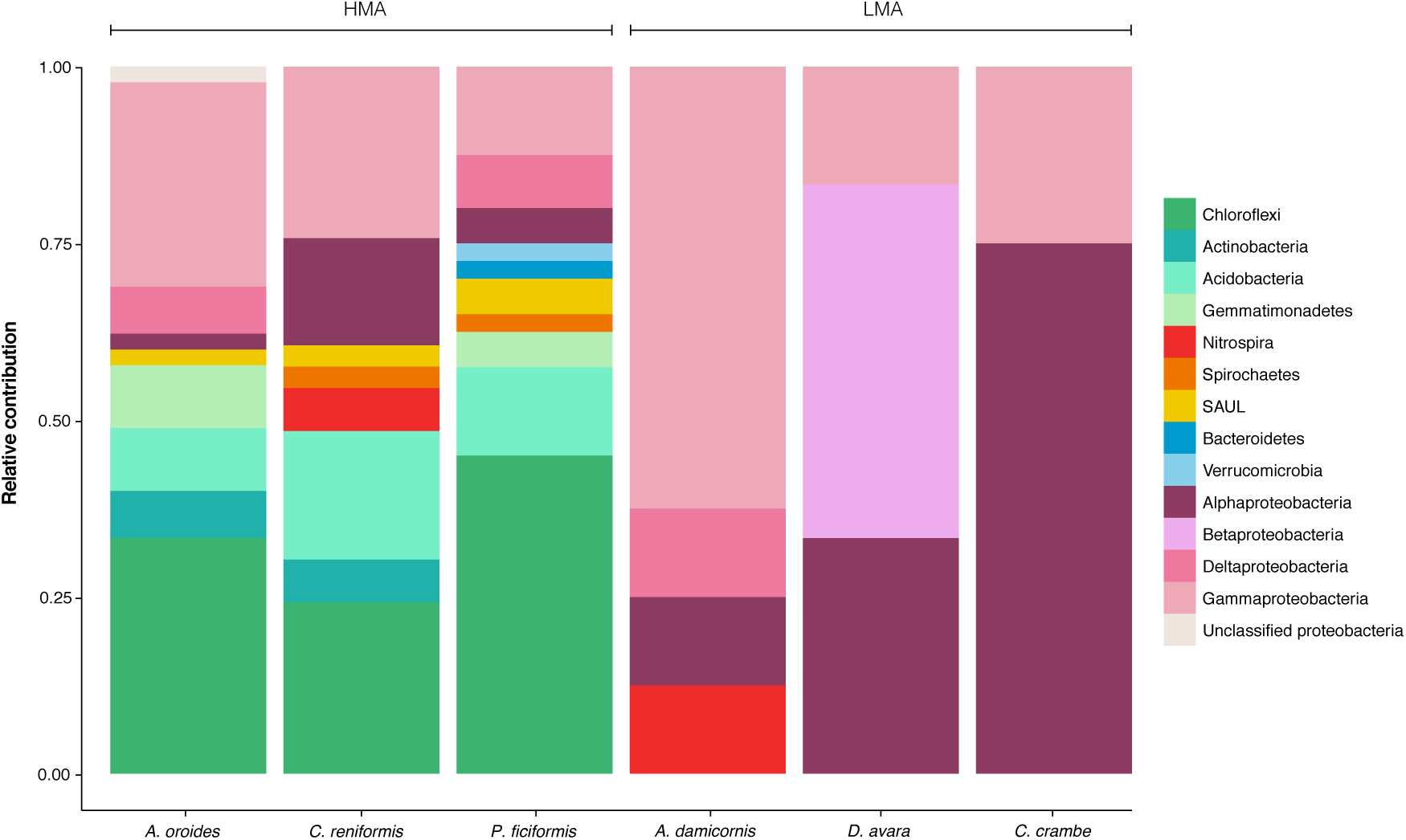
Taxonomic profiles of core microbiomes across host species and lifestyles (HMA vs. LMA). Taxonomic classification at the phylum level of core taxa and their relative contribution to diversity (species richness) for each core microbiome. The core microbiomes of HMA hosts harbored a larger taxonomic diversity than those of LMA hosts.

Sponges typically harbor certain taxa highly specific to the phylum *Porifera*. These “sponge-specific” 16S rRNA gene sequence clusters are monophyletic and span 14 known bacterial and archaeal phyla ([48, 49]). Taxa that fall into these sponge-specific clusters are only detected at very low abundances outside of sponge hosts, e.g., in the seawater and sediment ([27, 50, 25]). Some of these taxa are transmitted vertically from parent to offspring, suggesting sponge-microbe coevolution and cospeciation ([51]). We found that the core microbiomes of HMA hosts harbored a larger proportion of taxa that corresponded to sponge-specific clusters than the cores of LMA hosts (Table 2; Figure S4). Furthermore, these taxa had a higher average monthly abundance in the core microbiomes compared to the transient and opportunistic assemblages (Table S1). Then, despite the fact that identifying what constitutes a core microbiome remains elusive ([6, 25, 9]), the definition of the core used here (i.e., taxa present in at least 70% of the time series) provided a clear distinction in species composition between the core microbiomes and the transient and opportunistic assemblages (Figure2) that displayed prominent differences in the average monthly abundance of sponge-specific taxa (TableS1) and in their temporal turnover (Figures S1).

**Table 2:** Percentage of taxa within each assemblage and host assigning to sponge-specific clusters.

|  | HMA |  |  | LMA |  |  |
| --- | --- | --- | --- | --- | --- | --- |
|  | *A. oroides* | *C. reniformis* | *P. ficiformis* | *A. damicornis* | *D. avara* | *C. crambe* |
| Core | 42.2 | 45.6 | 60 | 25 | 0 | 12.5 |
| Transient | 43.3 | 40.7 | 47.1 | 25.8 | 9.1 | 9.8 |
| Opportunistic | 22.8 | 33.9 | 35.6 | 22.2 | 9.5 | 11.8 |

The innate immune defense of some sponge species can differentiate between pathogens, food bacteria and commensals in a manner similar to the adaptive immune system of vertebrates ([52, 53, 54, 55, 56, 57]). Thus, the above findings suggests that sponge-specific clusters, although present in the water column and sediment as part of the rare biosphere ([58, 50]), represent taxa important for host functioning.

### Core taxa are more stable and abundant

The unusual temporal coverage of our dataset allowed us to address a number of fundamental questions related to the temporal variability within and across microbiomes. We computed the mean abundance and the coefficient of variation 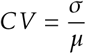 for each taxon’s abundance trajectory over time. What emerged for each microbiome was a negative relationship between the coefficient of variation and the log mean abundance: individual core and transient taxa were more stable and abundant than opportunistic taxa. However, in a few microbiomes, abundant conditionally rare taxa (CRTs) greatly outnumbered the abundance of core taxa. CRTs are extremely rare or below detection limit throughout most of the time series, but occasionally reach high abundance ([59]). Our results suggest that the presence of these taxa can have a negative impact on the stability of the focal core microbiome (Figure4 AB; FigureS5).

**Figure 4:**
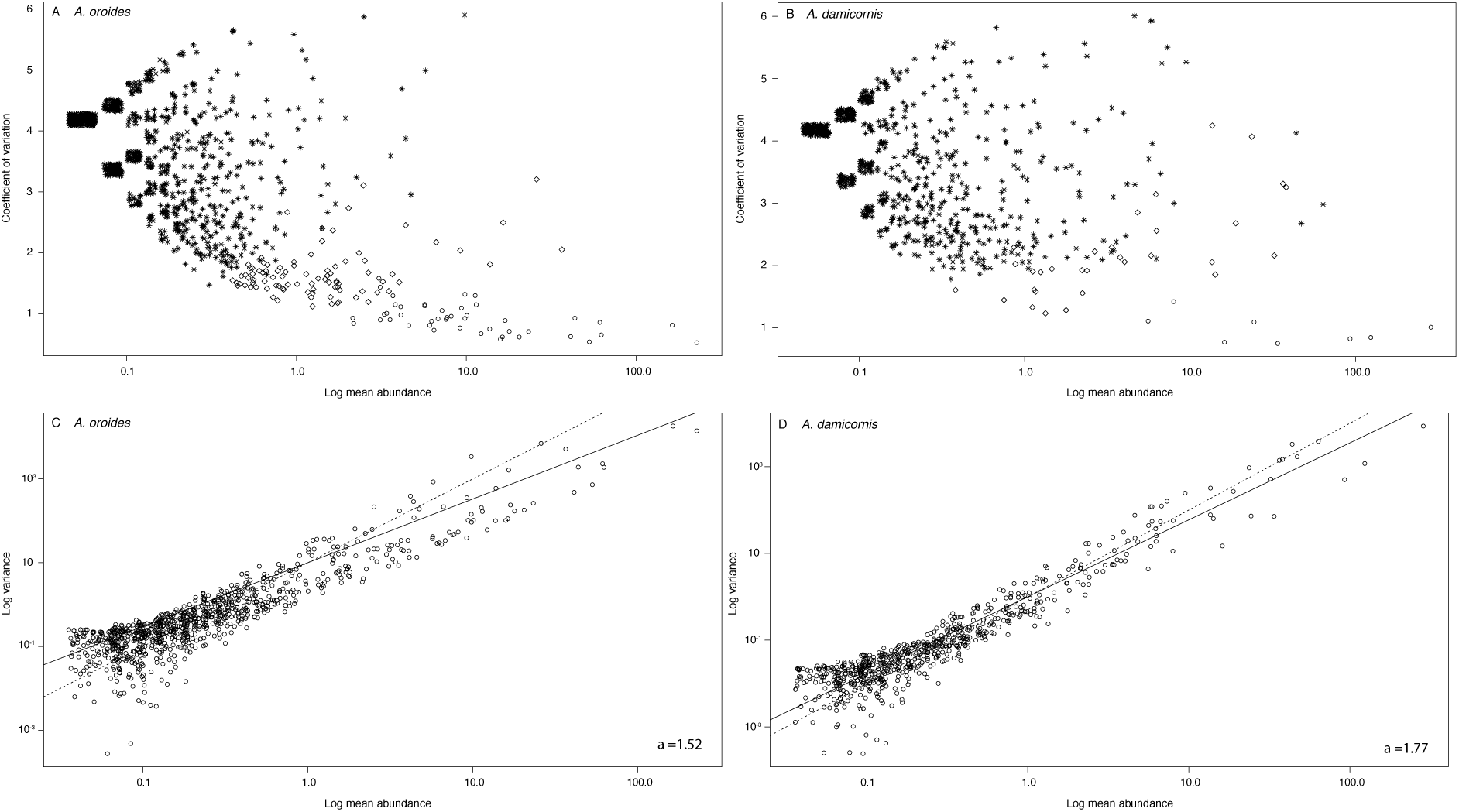
The top panel shows the relationship between temporal variability (CV) and mean log abundance for each taxon’s abundance trajectory over time, for host (A) *A. oroides* and (B) *A. damicornis*. Overlaying points have been separated jitter (random noise). Opportunistic, transient and core taxa are represented by stars, diamonds and circles, respectively. Individual core and transient taxa are more stable (Kruskal-Wallis test: *H* =2198, df=2, P<0.001 two-tailed; Dunn’s post-hoc test with bonferroni correction; P <0.001) and abundant (Kruskal-Wallis test: *H* =1694, df= 2, P<0.001; Dunn’s post-hoc test with bonferroni correction; P< 0.001 two-tailed) than opportunistic taxa. The bottom panel shows the relationship between log variance and log mean abundance for each taxon’s abundance trajectory over time (Taylor’s power law), shown for host (C) *A. oroides* and (D) *A. damicornis*. Solid lines represent the null expectation of Taylor’s power law, i.e., an exponent *a*=2, and the dashed lines correspond to the exponent of each host’s microbiome: 1.52 for *A. oroides* and 1.77 for *A. damicornis*.

There are several non-mutually exclusive mechanisms that can underpin the observed relationship between temporal variability and mean abundance. First, in agreement with recent observations on a broad range of marine communities, neutrality alone is unlikely to explain the high abundance of the most common species ([60]). Second, lower variability can imply the presence of self-regulation (i.e., density dependent processes) at higher population abundances ([61, 62]). Furthermore, on a log-log scale, our observed relationship between temporal variability (now variance *σ*^2^) and mean abundance describes Taylor’s power law ([63]). The null expectation of the temporal version of Taylor’s law states that the variance scales with the mean abundance following a power law with an exponent equal to 2. This implies that population or community variability is constant. Conversely, if the exponent is less than 2, then the variability in population abundance decreases with increasing mean population abundance, as seen in Figure 4 CD and Figure S6. Third, for large population sizes, as is characteristic of microbial communities like those studied here, environmental stochasticity is more likely than demographic stochasticity to reduce Taylor’s power law exponent ([64, 65]). Fourth, even weak interactions among species can reduce this exponent; for example, interspecific interactions, such as competition typically lead to smaller fluctuations of common versus rare species within communities ([65])-a result that could be extended to other types of interactions. Below, we explore the contribution of these mechanism to microbiome dynamics and stability in more detail.

### Core microbiome density, not diversity, begets its stability

We aggregated all individual microbial population abundances within each temporal assemblage–core, transient, and opportunistic (Figure 5). This revealed two markedly different temporal dynamics across microbiomes: in the hosts *A. oroides, C. reniformis* and *C. crambe*, core microbiomes were very dense, i.e., they accounted for the majority of microbiome relative abundance. In contrast, the core microbiomes of hosts *D. avara, A. damicornis* and *P. ficiformis* were sparser, and instead transient and/or opportunistic assemblages dominated microbiome relative abundance. We found that dense cores were more stable over time than sparse cores, measured as community-level invariability (the inverse of variability, [66]; Figure 5). The association between core density and stability was also robust to a more stringent definition (≥85%) of the core microbiome (Figure S6). For some microbiomes, no taxa persisted on or above this threshold.

**Figure 5:**
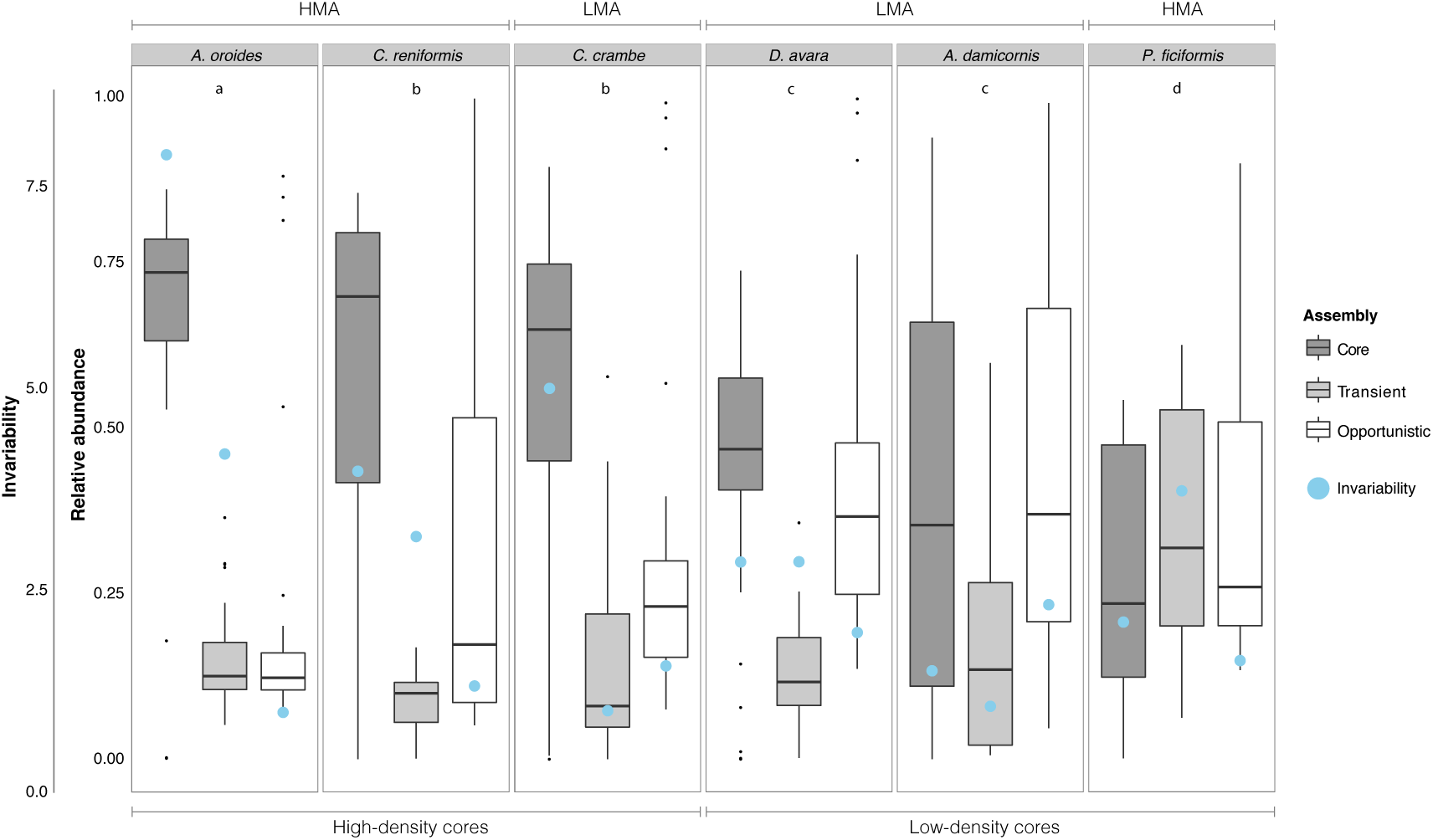
The contribution of each assemblage to microbiome abundance and its aggregated stability across hosts. The inner y-axis shows the contribution to microbiome relative abundance by each assemblage across host species. Each box shows the median including the first and third quartiles (the 25*^th^* and 75*^th^* percentiles), representing temporal variation. The outer y-axis shows invariability at the community level (blue dots) for each assemblage. The figure is ordered from the highest to the lowest in terms of core microbiome density. Lowercase letters denote different significant scenarios (see Table S2 for more detailed information): (a) The core microbiome was significantly different from the transient and opportunistic assemblages, but transient and opportunistic assemblages were not significantly different from each other; (b) All assemblages were significantly different; (c) The core microbiome and the opportunistic assemblage were not significantly different, but the core microbiome and transient assemblage, and the transient and the opportunistic assemblage were significantly different from each other; and finally (d) No significant differences between any of the assemblages. What emerged was three high-density (scenario a & b), and three low-density (scenario c & d) cores, respectively.

Community-level stability showed a weak (non-significant) relationship with diversity (species richness; Figure S8 A). Instead, we observed a significant positive relationship between community-level stability and the median relative abundance (Figure S8 B). This agrees with some studies that demonstrated a positive relationship between community-level stability and the relative abundance or biomass of common plant species ([67, 68, 69]). Importantly, this suggests that the effect of diversity on stability can be constrained by the dynamics of only a few common and abundant species, as seen in the case of our high-density cores (Figure 5); for example, host species *P. ficiformis* harbored the second largest core diversity (*S*=40, Table 1) that still resulted in the least stable low-density core. On the other hand, sponge species *C. crambe* achieved a stable high-density core by harboring only a few core taxa (*S*=8, Table 1).

Furthermore, in agreement with the HMA-LMA dichotomy, the metabolic profiles of *P. ficiformis* and *C. crambe* yet match those of other archetypal HMA and LMA hosts, respectively ([70]). While *P. ficiformis* harbored an unstable low-density core, its core was nonetheless comprised of the largest proportion of sponge-specific clusters (Table 2; Figure S4), suggesting that microbiome stability may not always be a prerequisite for host functioning.

### Intraspecific interactions drive core microbiome dynamics

In order to further disentangle the main drivers of core microbiome temporal dynamics, we used a model that decomposes temporal fluctuations in species abundances into three contributions–Interspecific interactions, intraspecific interactions, and environmental variability. Environmental variability includes apart from ecological drift, the aggregated effects of the host on the microbial consortium, as well as the external environment acting on the host. While we included several environmental covariates that had been recorded over the 36 months, such as water temperature, salinity and nutrients, our model is able to capture unmeasured effects through the use of latent variables (see *Methods* for more information).

We found that intraspecific interactions explained the largest proportion of variation, suggesting that all core taxa on average, experienced strong self-regulation (Figure 6 A). There was a marked difference between HMA and LMA host species in terms of the variation explained by interspecific interactions: In the core microbiomes of the HMA hosts, interspecific interactions had a relatively large effect on the dynamics, whilst this effect was almost negligible in the LMA host species (Figure 6 A).

**Figure 6:**
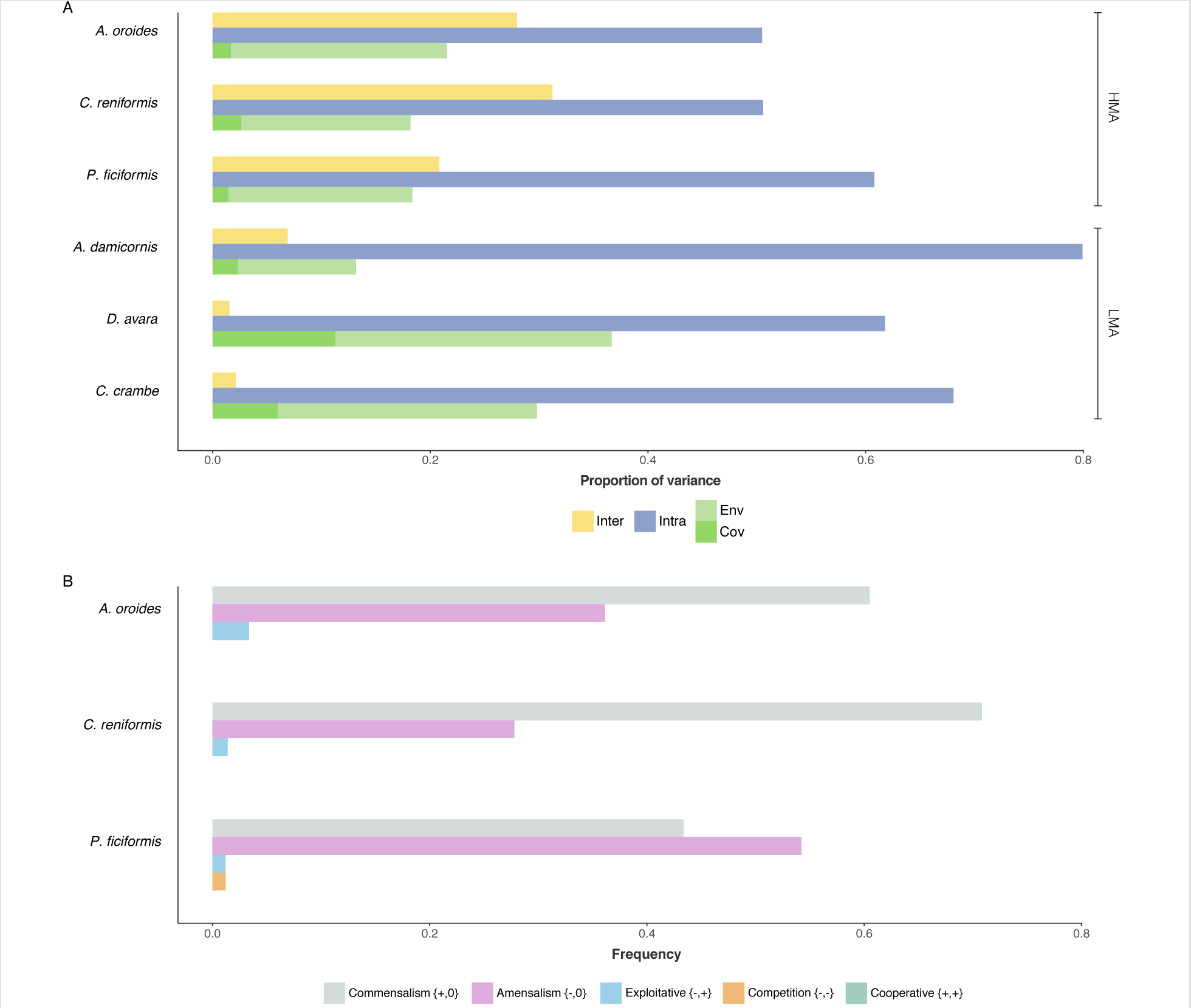
Processes explaining core microbiome dynamics and the frequency of of all possible interaction types within HMA core microbiomes. Panel A shows the relative contribution of interspecific (yellow) and intraspecific (blue) interactions and environmental variability (greens) to temporal variation in microbial population abundances across core microbiomes. Environmental variability is decomposed into residual variation (light green, Env) and variation attributed to the modeled environmentalcovariates (dark green, Cov). In all hosts, core microbiome dynamics were mainly driven by intraspecific interactions. While the modeled environmental covariates explained relatively little variation across core microibomes, an important driver of the dynamics in HMA cores was interspecific interactions which were almost negligible in LMA cores. Panel B shows the relative frequency of all inferred possible interaction types within HMA cores. This was calculated from the marginal sign posterior distribution of *alpha_i,j_* (see *Methods*). Commensalism {+,0} and amensalism {-,0} were the most frequent interaction types across HMA cores. Competitive {-,-} and exploitative {+,-} interactions were exceptionally rare. Noteworthy, cooperative interactions {+,+} was never inferred.

The main drivers we identified for core microbiome temporal dynamics somewhat differ from those commonly reported for other large species-rich systems; for example, studies assuming Lotka-Volterra dynamics in macroecological communities ([71, 72, 73, 74]) or other types of time series analyses applied on free-living marine microbial communities ([75, 76, 77]) often report environmental variability as the major factor affecting population dynamics. While environmental variability was an important factor, intraspecific interactions was the single most important driver affecting the dynamics across all core microbiomes. Furthermore, the modeled environmental covariates only accounted for a small fraction of the variation explained by environmental variability (Figure 6 A), indicating that the sponge microbiome may experience a reduced influence of the external environment acting on the host.

A large body of theoretical and empirical literature suggests that self-regulation, which we equate here to intraspecific interactions, is a key determinant of temporal stability ([61, 62]). The simple premise of this regulatory mechanisms is that species abundances decrease per capita growth rates when population abundances are high and vice versa. Recently *Barabàs and colleagues* (2017) showed that stability requires the majority of species in a community to experience negative self-effects. Although this work focuses on asymptotic stability (i.e., whether small perturbations of species’ abundances away from an equilibrium point tend to be dampened, with the system returning to equilibrium) and not on temporal variability, their results in combination with ours suggest that strong self-regulation should also be the norm within microbiomes ([78]).

### The core microbiomes are comprised of weak unilateral interactions

The nature and strength of interspecific interactions are pivotal to understand the dynamics of species interaction networks ([79, 80]). Our unique temporal series allows for determining how microbes interact with each other within different host species. Because interspecific interactions were almost negligible in the LMA hosts, we analyzed the most credible network structure of each HMA core microbiome (Figure 7; Figures S9). For these three networks, only a small fraction among the possible interactions were likely to occur (Figures S10), resulting in low network connectance (5-7%). The networks had a skewed distribution of interaction strengths toward many weak and a few strong interactions (Figures S11). This pattern mimics that found in empirical food-webs [81, 82]) and mutualistic networks ([83]). Theory shows that skewed distributions of interaction strengths beget stability, and arises during the assembly of persistent communities ([84, 85]).

**Figure 7:**
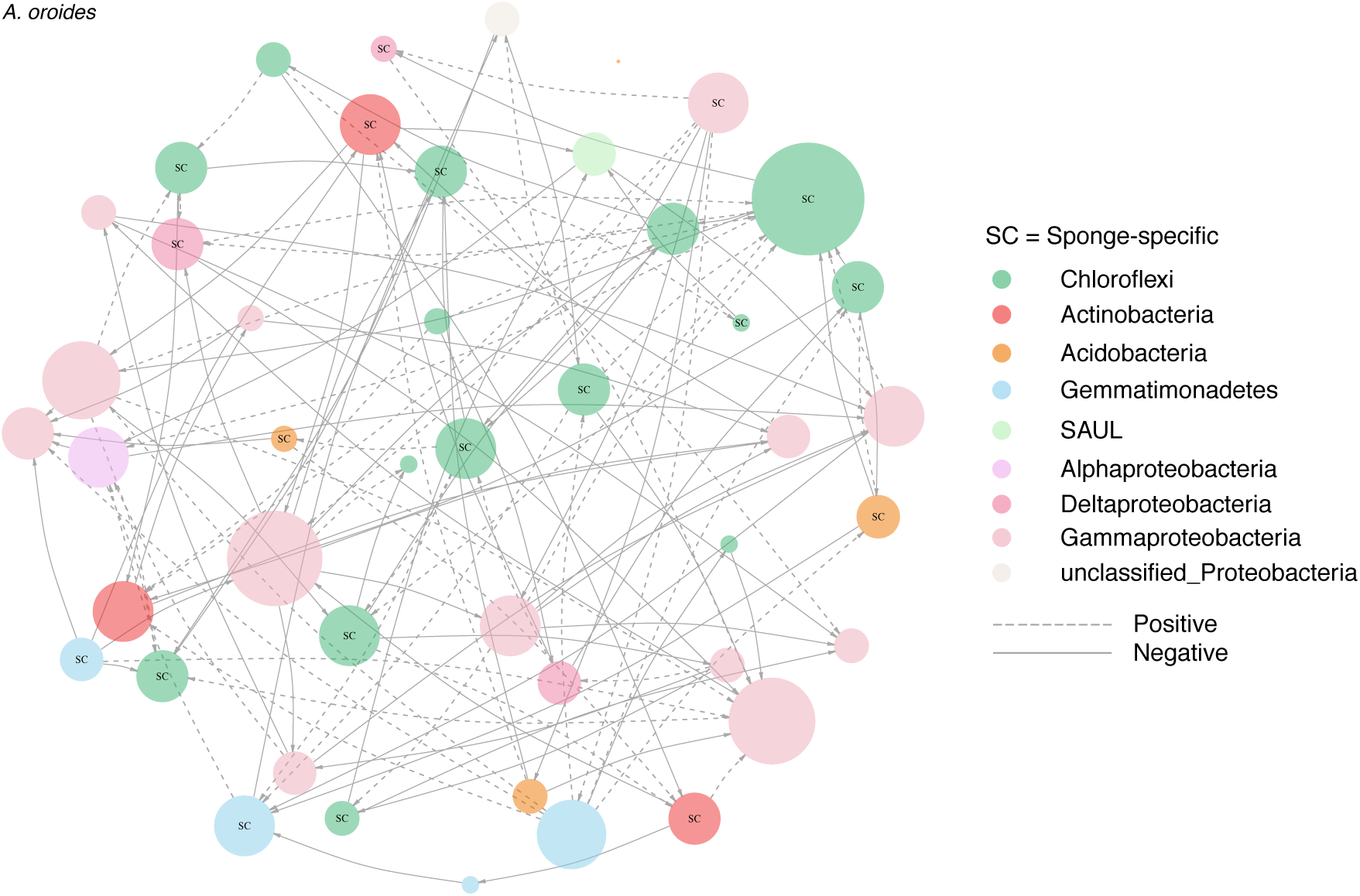
The most credible network structure for HMA host *A. oroides*’s core microbiome. Nodes represent core taxa and links their inferred ecological interactions. Node size is scaled to their degree (i.e. in and out-going links). Colors correspond to different bacterial phyla and dash and solid lines represent positive and negative interactions, respectively. Nodes marked with SC correspond to taxa that assigned to sponge-specific clusters. See Figure S10 for the corresponding link probabilities between OTU *i* and *j*, and Figure S9 for the networks belonging to the core microbiomes of HMA hosts *C. reniformis* and *P. ficiformis*. See *Methods* for how the networks were constructed.

Theory shows that reciprocal interactions such as exploitation {+/-}, cooperation {+/+} and competition {-/-} differ in their effects on community stability and ecosystem functions ([86, 87, 88, 89, 90, 79, 80]). Specifically, communities consisting of a mixture of unilateral interactions are more stable than those with only reciprocal interactions ([80]). We found that amongst all possible interspecific interactions (Figures S12), unilateral interactions in the form of commensalism {+,0} and amensalism {-,0} dominated (Figure 6 B). Reciprocal interactions–cooperative {+,+}, competitive {-,-} and exploitative {+,-} interactions were exceptionally rare. The core microbiomes of hosts *A. oroides* and *C. reniformis* were largely dominated by commensal interactions, whereas the core microbiome of host *P. ficiformis* had a higher frequency of amensalism. It was also the only core network that had competitive interactions among its members, although at very low frequencies. Sponge-specific clusters, although prevalent in the core microbiomes of HMA hosts were not more connected than other nodes within these networks

Certainly, opportunistic and transient taxa can influence the dynamics of the core microbiome. However, we focused on inferring interspecific interactions among core taxa for three main reasons: (1) Computation–Including non-core taxa would increase network size ten folds, and thus not be computationally feasible; (2) Information–There is very little information in taxa only occurring in one or a few timesteps, thus the inference for those taxa would be very poor; and (3) Ecology–Interspecific interactions require species to frequently co-occur. The many occasional taxa observed throughout the time series, especially those occurring at high densities (i.e., the CRTs) are likely acting together with ecological drift as sources of stochastic variability. In our model, the effects of unmodeled interspecific interactions are captured by the inclusion of the latent variables (see *Methods* for more information).

### Microbial transmission mode can affect microbiome stability and functioning

High-density cores (i.e., those core microbiomes whose taxa accounted for the majority of abundance) were found in sponge species that transmit microbes vertically from adult to larvae ([91, 92, 93]), whereas low-density cores represent sponge species with larvae largely deprived of microbes ([94, 95, 96]).

Vertical transmission provides an evolutionary mechanism for preserving particular combinations of microbes, including their interaction structure and the ecosystem functions that emerge from it ([97]). Evidence from other ecological communities, including the human gut microbiome, suggest that priority effects–the order and timing of species arrivals–determine the interactions among species, and in turn also community assembly and stability ([98, 99, 100]). The process of vertical inheritance of microbes likely has similar outcomes as priority effects. Furthermore, as pathogenic microbes use cooperative secretions that modify their environment to enhance their growth and expansion ([101, 102]), it is reasonable to assume that commensal microbes do too. We therefore hypothesize that the complementary set of microbes that are vertically transmitted from parent to offspring pre-empt the initial host niche by fast reaching carrying capacity, while simultaneously modifying it in their favor. This would inhibit the subsequent colonization of some microbes, while facilitating the establishment of certain others.

As previously mentioned, host species *P. ficiformis* harbored an unstable low-density core, yet it harbored the largest consortia of sponge-specific clusters within its core. This does not only suggest that microbiome stability may not always be a requirement for host functioning, it also indicates that host species *P. ficiformis* achieves HMA archetypal functional characteristics by means of horizontally selecting commensal microbes from the water column, many of which fall into sponge-specific clusters. These sponge-specific microbes are examples of conditionally rare taxa (CRTs) outside the sponge host, but through some recognition mechanism are allowed to flourish within the host.

## Conclusion

Natural systems commonly work as temporal networks ([103]). To increase our understanding of the processes governing microbiome assembly, stability and functioning, this study highlights the importance of defining the core microbiome longitudinally (i.e., analysing temporal dynamics) rather than cross-sectionally. Most studies to date have focused on the human microbiome (see e.g., [104, 13, 105, 79, 100]), with only a few studies exploring temporal dynamics in other host systems ([11, 106, 107]). By focusing on the six most common sponge species of the temperate Mediterranean benthic community, we successfully characterized microbiome dynamics and stability under natural conditions.

The observed negative relationship between temporal variability and abundance that emerged from the analyses of each microbiome is conducive to the notion of the dynamic core microbiome. This notion allowed us to determine that irrespective of host’s eco-evolutionary characteristics and lifestyles, it is the density of the core microbiome rather than its diversity that determines the stability of the sponge microbiome. We hypothesize that priority effects mediated by vertical transmission underpins this pattern, which may further suggest that high-density cores confer hosts resistance against the establishment of occasional taxa that sponges are constantly exposed to through their filter-feeding activities. The core microbiome has been proposed as the common taxa shared among microbiomes in an habitat ([108, 6]). Our finding reveals a relevant functional attribute that constitutes an step forward towards the identification and characterization of the so-called core microbiome.

We further found that intraspecific self-regulation is much more important for microbiome dynamics than environmental forcing. The most credible core interaction networks consisted of members interacting weakly with each other with a dominance of comensal and amensal interactions. Altogether, this suggests that host-associated microbiome dynamics and its emerging interaction structures differ from the temporal dynamics of free-living microbial communities. These interactions have likely coevolved to maintain host functionality and fitness over ecological, and even evolutionary time scales.

We have focused on compositional stability over time–a notion inclusive of variation in both species relative abundances and species composition. More generally, compositional and functional stability often show complex interdependencies ([109]), because, for example, compensatory dynamics between species may reduce functional loss within communities ([110]). For most host species studied here, our results suggest that there may exist a positive relationship between compositional and functional stability. However, for host-associated microbiomes in general, it is difficult to disentangle the effects of compositional stability from the host’s own ability to control the identity and abundance of its microbes on the overall functioning. For one pair of hosts, our findings of high-density and low-density cores did not match the notion of a positive relationship between compositional and functional stability. This suggests that, at least for some host species, functioning is achieved by other mechanisms that do not require compositional stability. The relationship between microbiome compositional stability and host functionality will benefit from further investigation that focuses on the temporal dynamics and functioning of the core microbiome.

## Methods

### Sponge collection

The sponge species *Agelas oroides, Chondrosia reniformis, Petrosia ficiformis, Axinella damicornis, Dysidea avara* and *Crambe crambe* were collected monthly from March 2009 until February 2012 close to the Islas Medas marine reserve in the NW Mediterranean Sea 42°3′0″*N*, 3°13′0″*E* by SCUBA at depths between 10-15 metres. Three replicates per sponge species (i.e., different individuals per sampling time) and ambient seawater samples were collected. The collected sponge species belong to six different orders that represent common members of the Mediterranean benthic community. Each sponge species were identified based on distinct morphological features. Specimens were sublethally sampled and excised fragments were placed in separate plastic bottles and brought to the surface were they were frozen in liquid nitrogen until DNA extractions. Samples of the ambient water were taken at 5 m depth and poured into three separate 5 L jars. All samples were stored at −80°C until DNA extraction. Aliquots of seawater (300-500 mL each, 1 aliquot per sample jar) were concentrated on 0.2 *µ*m polycarbonate filters, submerged in lysis buffer and stored at −80°C until DNA extraction.

### DNA extraction and sequencing

Following the manufacturers Animal Tissue protocol, 16S rRNA gene sequences were PCR-amplified from sponge samples (n=648) and seawater filters (n=108) using the DNeasy tissue kit (Qiagen, CA, USA), and subsequently submitted to the Research and Testing Laboratory (Lubbock, TX, USA) for gene amplicon pyrosequencing. Samples were amplified with primer 28F and amplicons were sequenced using 454 Titanium chemistry (Roche, CT, USA), producing 250 base pair read lengths in the 5′→3′ direction.

### Analysis of sequencing data

454 reads were processed in mothur v.1.29.2 ([111]). Raw reads were pooled from replicates belonging to the same sponge species. Fasta, qual and flow files were extracted from binary sff files; sffinfo(*…*,flow=T). Flow files were then filtered based on barcodes to speed-up the proceeding de-noising process; trim.flow. Sequences were de-noised; shhh.flows(*…*, lookup= LookUp_Titanium.pat). The LookUp-file is necessary and specific to the 454 technology used. Next the barcode and primer sequences were removed together with sequences shorter than 200bp and/or contained homopolymers longer than 8bp; trim.seqs(*…*, pdiffs =2, bdiffs =1, maxhomop =8, minlength =200). In order to minimise computational effort, files were reduced to non identical sequences; unique.seqs. Non redundant sequences were aligned to SILVA 102 reference alignment with default kmer search and Needleman-Wunsch algorithm; align.seqs(*…*, flip =F). Non overlapping sequences were removed; screen.seqs(*…*, optimize= end, start= 1044, criteria = 95), in addition to empty columns that were introduced from the alignment process; filter.seqs(*…*,vertical =T, trump =.). Aligned sequences were reduced to non redundant sequences; unique.seqs. To further reduce amplification errors, less abundant sequences were binned to more abundant sequences if they were within 2bp of a difference; pre.cluster(*…*, diffs =2). Chimeric sequences were identified; chimera.uchime(*…*, dereplicate =T) and removed; remove.seqs. Sequences were classified using the RDP reference taxonomy; classify.seqs(*…*, template =trainset9_ 032012.pds.fasta, taxonomy =trainset9_ 032012.pds.tax, cutoff =80), and non bacterial lineages were removed; remove.lineage(*…*, taxon= Mitochondria-Chloroplast-Archaea-Eukaryota-unknown). We calculated pairwise distances between aligned sequences; dist.seqs(*…*, cutoff =0.050).

Due to an uneven sequence distribution across samples, we pooled sequences across the three host species replicates prior to OTU clustering. As we found a plate effect on the number of sequences, we sub-sampled 1500 sequences from the resulting monthly samples. This number corresponded to the average of the three lowest host-plate averages. Sequences were thereafter clustered into OTUs defined at 97% similarity; classify.otu(*…*, label=0.030) and outputted to an OTU-table (.shared-file); make.shared(*…*, label=0.030).

### Identification of sponge-specific clusters

A representative sequence from each OTU was taxonomically assigned using a BLAST 62 search against a curated ARB-SILVA database containing 178 previously identified sponge-specific clusters (SC) ([49]). For each BLAST search, the 10 best hits were aligned to determine sequence similarities. The most similar OTU sequence to the respective reference sequence within the database was then assigned to an SC based on a 75% similarity threshold: (i) a sequence was only assigned to any given SC if its similarity was higher to the members of the cluster than to sequences outside the cluster; and (ii) if its similarity to the most similar sequence within the cluster was above 75%. A majority rule was applied in cases where the assignment of the most similar sequences was inconsistent, and the OTU sequence was only assigned to the SC if at least 60% of the reference sequences were affiliated with the cluster.

### Microbiome stability and dynamics

#### Temporal turnover

We applied a measure of temporal turnover that describes the extent to which individual OTUs and consequently the microbiome changes over time ([47]. Importantly, this measure decomposes abundance fluctuations into two additive contributions of change due to microbiome composition and total abundance.

Total turnover *D* between times *t* and *u*, (*u* > *t*) is defined as

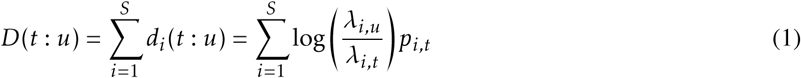

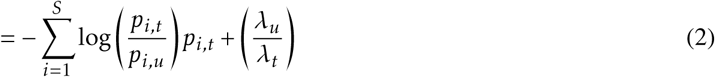

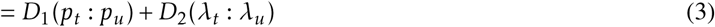

where 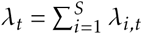 represent the sum of the expected total abundance of each OTU in the microbiome. The expected abundance of OTU *i* in time *t*, i.e., *λ_i,t_,i* = 1,2,…,*S* is unknown and therefore needs to be estimated from an observed time series. *p_i,t_* represents the relative abundance of OTU *i* in time *t*, and is calculated as 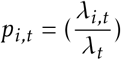. As such, the total turnover *D* can be decomposed into *D*_1_ which is related to the amount of change in microbiome composition, and *D*_2_ reflecting the amount of change in total abundance.

As noted above, the expected abundance needs to be estimated. We thus modeled each time series of time series of high-throughput DNA sequence *N_i,t_* assuming a Poisson log-linear model with a time-varying mean parameter *λ_i,t_*

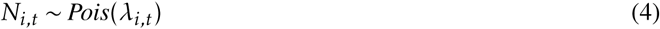

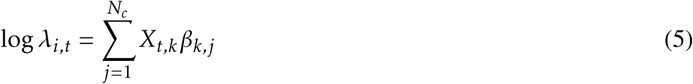

where *X_k,t_* is a time series of *k* = 1,2,…,*N_c_* environmental covariates, and *β_k,j_* is the corresponding regression coefficient that needs to be estimated. We included temperature, salinity, chlorophyll, bacterial cell density, nitrite (*NO*_2_), ammonia (*NH*_4_), and phosphate (*PO*_4_) as the *N_c_* environmental covariates. All covariates where standardized to have zero-mean and unit variance.

#### Community-level stability

We applied a newly developed measures of temporal stability defined as invariability ([66]) at the community-level, i.e., for each core microbiome, transient and opportunistic assemblages, respectively. Invariability at the community-level is defined as

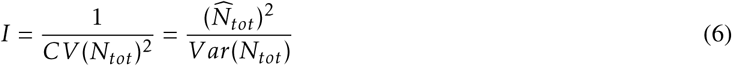

where 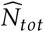 and 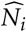 denote the average of the total abundance and the average of the abundance of OTU *i* over the time series, respectively.

#### Core dynamics and ecological interactions

We adapted a multivariate first-order autoregressive model assuming Gompertz population dynamics ([112]) to model core dynamics and to infer ecological interactions among core taxa from time series of high-throughput DNA sequence counts. Compared to the commonly used Generalized Lotka-Volterra (GLV) model (see e.g., [113]), the Gompertz model is linear on the log-scale and thus a good approximation of nonlinear dynamics. Furthermore, a severe problem with even the simplest GLV model is that the number of parameters that need to be estimated often is larger than the number of available data points. **Mutshinda and colleagues* (2009)* addressed this problem by using Gibbs Variable Selection ([114]) in order to induce sparseness: the *N(N* + 1) interspecific interactions are constrained so that most are shrunk to zero. We further adapted this framework to address two additional challenges of microbiome data. First, to model covariances between a large number of taxa using a standard multivariate random effect is computationally challenging: the number of parameters that has to be estimated when assuming a completely unstructured covariance matrix increases quadratically as the number of taxa increases. We therefore parameterized the residual covariance matrix using a latent variable approach. Latent variables also capture unmeasured effects (e.g., host effects and unmodeled interspecific interactions) which if not accounted for may lead to erroneous inference ([115]). Second, high-throughput DNA sequencing produces compositional data, i.e., non-negative counts with an arbitrary sum imposed by the sequencing platform, which can produce spurious correlations if not properly accounted for (see e.g., [116, 117, 118]). We therefore used a log-linear model with a Poisson likelihood that includes a random effect accounting for sample size. This model produces a likelihood equivalent to that of the logistic normal multinomial model but is more convenient for computation and estimation ([119]).

#### Process model

To accommodate the time series, we refer to OTU *i* ∈ {1,…,*I*} in time point *t* ∈ {1,…,*T*}. Let *N*(*µ,σ*^2^) denote a univariate normal distribution with mean *µ* and variance *σ*^2^, and analogously, let *MVN*(*µ,Σ*) denote a multivariate normal distribution with mean vector and covariance matrix *Σ*. If we denote *n_i,t_*\* as the expectation of *n_i,t_* which is the natural logarithm of the observed time series *N_i,t_*, then on the natural logarithmic scale we have the expected number of 16S rRNA gene sequences from OTU *i* in time point *t* within a given core microbiome described by

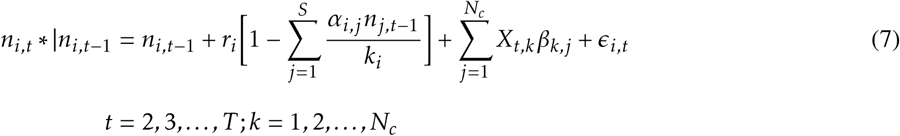

where we assume *r_i_* ~ N (0,10) and *K_i_* ~ Exp(1). The coefficients measuring each taxon’s response the *k* - *th* environmental covariate are assumed *β_k,j_* ~ N (0,100). We assumed correlated residual responses to the environment by included temperature, salinity, chlorophyll, bacterial cell density, nitrite (*NO*_2_), ammonia (*NH*_4_) and phosphate (*PO*_4_) as the *N_c_* environmental covariates potentially influencing the modeled core OTUs. All covariates where standardized to have zero-mean and unit variance. *ϵ_i,t_* ~ MVN (0,Σ) represents the residual variance, where Σ is the residual covariance matrix which was parameterized using latent variables.

#### Latent variables

As previously stated, to improve statistical efficiency we parameterized the residual covariance matrix Σ using a latent variables approach, where the index *q* runs over the *q* = 1,…,2 latent variables.

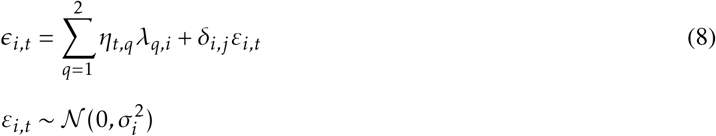

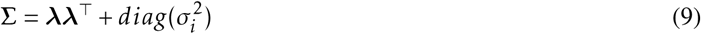

were *η* denote the latent variables and λ their corresponding factor loadings. Both are assigned standard normal priors *N* (0,1) with the assumption of zero mean and unit variance to fix the location (see chapter 5, [120]). *δ_i,j_* denote the Kronecker’s delta such that *δ_i,i_* = 1 and *δ_i,j_* = 0 for *i* ≠ *j*. Thus, the covariance matrix Σ can be computed from the factor loadings (Equation 9). The diagonal elements of the covariance matrix Σ quantify the amount of residual variation for OTU *i* not captured by the modeled environmental covariates.

#### Observation model

The time series of high-throughput DNA sequence counts *y_i,t_* were modeled as Poisson random variables with means *λ_t,i_* satisfying the following log-linear model

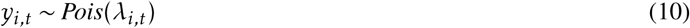

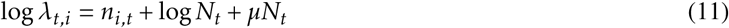

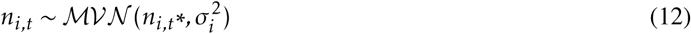

where log*N_t_* is an offset accounting for the observed total number of DNA sequences in time *t*, while *µN_t_* ~ *N* (0,100) represent the random effect accounting for sample size in time *t*. Both thus represent the total abundance in time *t*, and account for the aforementioned compositional nature of high-throughput DNA sequencing data.

#### Gibbs Variable Selection

As mention above, we used Gibbs Variable Selection ([114]) method in order to constrain the model to only use interspecific interaction coefficients *α_i,j_* for which there were strong support in the data. This was achieved by introducing a binary indicator variable *γ_i,j_* for *i* ≠ *j*, and assuming *γ_i,j_* ~ *Bernoulli*(*p*), such that *γ_i,j_* = 1 when OTU *j* is included in the dynamics of OTU *i*, and *γ_i,j_* = 0 otherwise. Where there was low support for *α_i,j_* in the data, *γ_i,j_* = 0 and the corresponding interaction was excluded from the model. On the other hand, when *γ_i,j_* = 1, *α_i,j_* was freely estimated from the data. The parameter *p* represents our prior belief about the proportion of realized interspecific interactions: we set *p*=0.1, thus assuming that 90% of all interspecific interactions were zero.

We used Markov chain Monte Carlo (MCMC) simulation methods through JAGS ([121]) in `R` ([122]) using the *runjags* package ([123]) to sample from the joint posterior distribution of all the model parameters. We ran 10 independent chains with dispersed initial values for 5*e*6 iterations, discarding the first 2*e*6 samples of each chain as burn-in and thinned the remainder to every 50*th* sample. We evaluated convergence of model parameters by visually inspecting trace and density plots using the packages *coda* ([124]) and *mcmcplots* ([125]), as well as using the Geweke diagnostic ([126]). In addition, to ensure good mixing of the parameter *α_i,j_*, we calculated the number of jumps the parameter *γ_i,j_* made between its two states, 0 and 1.

#### Variance partitioning

The total variance *V_i_* affecting the dynamics of core OTU *i* can be decomposed into additive sources reflecting interspecific interactions, intraspecific interactions, and environmental variability (i.e., measured environmental covariates and residual variation) as follows,

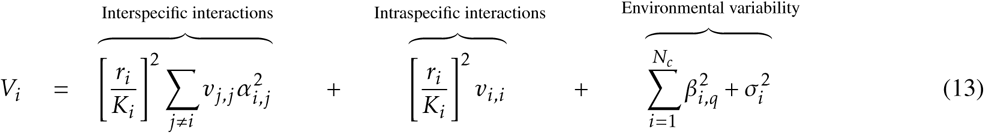

where *v_i,i_* represent the stationary variance for *n_i_* (Equation 11), 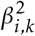 the variance attributable to each *k* covariate, and 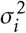 correspond to the residual variance (the diagonal elements of Σ, Equation 9). As a consequence of Equation 13, the proportion of variation attributed to e.g., interspecific interactions can be calculated as

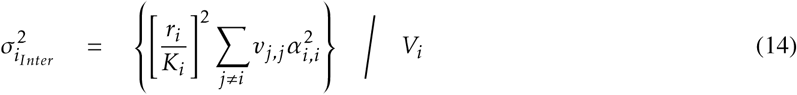

#### Finding the most credible network structure

To characterize each core microbiome network, we analyzed the interaction structure of the marginal posterior distribution for the interaction coefficient *α_i,j_*. The parameter *α_i,j_* is a probability distribution, thus containing the probability of OTU *j* having a per capita effect on the growth rate of OTU *i*. This means that we can decompose *α_i,j_* into two marginal posterior distributions–one for interaction sign and another one for interaction strength. We constructed the core microbiome networks for each HMA host as means of visualizing the most credible network structures. This was done by mapping the marginal posterior distribution of the average number of links onto *α_i,j_*, thus extracting the marginal posterior average number of links with the highest probability of non-zero interactions. As a way of further validating network structure, we compared the marginal posterior distribution of connectance to the empirical connectance of each constructed network.

#### Code and Data Availability

All data and code will eventually be available at the Open Science Framework.

## Acknowledgements

JRB. was supported by an FPI Fellowship from the Spanish Government (BES-2011-049043), JMM. was supported by the French Laboratory of Excellence Project ‘TULIP’ (ANR-10-LABX-41; ANR-11-IDEX-002-02) and by a Region Midi-Pyrenees Project (CNRS 121090) and RC and MR were supported by a Spanish Government Project (CGL2013-43106-R) and by a grant from the Catalan Government (2014SGR1029).

## Author contributions

JRB and JMM conceived the study. JRB performed all the analyses and wrote the manuscript. JRB and RBO adapted the MAR(1) model. JRB and JMM refined the manuscript. RC and MR collected the data. All authors commented on later versions of the manuscript.

